# Assessment of *Agaricus bisporus* Mushroom as Protective Agent Against Ultraviolet Exposure

**DOI:** 10.1101/2021.10.21.465111

**Authors:** Chae Yeon Hwang, Yuniwaty Halim, Marcelia Sugata, Dela Rosa, Sherlyn Putri Wijaya, Eden Steven

## Abstract

Mushrooms are versatile materials with applications including but not limited to food, cosmetics, and pharmaceutical industries. In this work, the potential of the common button mushroom, *Agaricus bisporus*, as a protective agent against ultraviolet exposure was assessed. The assessment was done by investigating the radical scavenging activity, sun protecting capability, and tyrosinase inhibiting properties of *Agaricus bisporus* ethanolic extract. The extraction was carried out using absolute ethanol as its solvent at low to room temperatures. The bioactive components of the ethanolic extract were analysed for its phenolic and flavonoid contents quantitatively, while other phytochemical agents were analysed qualitatively. The *Agaricus bisporus* ethanolic extract was found exhibit varying degree of activity in all of the assessment. We found low radical scavenging ability with %RSA IC50 of ∼5456 *μ*g/mL, low to moderate sun protecting factor of ∼5.355 at 5000 ppm concentration, and high tyrosinase inhibition property with IC50 of ∼2 *μ*g/mL. The high tyrosinase inhibition property was found to correlate with relatively high total phenolic content of ∼1143 mg GAE/100g for *Agaricus bisporus* and the presence of terpenoid in the ethanolic extract.

## Introduction

Research of NASA scientists have evidenced that ground ultraviolet radiation (UVR) reaching the Earth’s surface has increased significantly over the past few decades ^1^. UVR lies on the left of visible light in the electromagnetic spectrum, meaning its wavelengths are relatively shorter and they contain more energy. These short wavelengths of UVR enable UVR to penetrate the human skin ^2^. Moderate exposure to UVR is actually required by humans to promote conversion of the 7-dehydrocholesterol into provitamin D and then into its functional form of vitamin D. However, excessive amounts of UVR has been proven to induce negative effects on human health as it may trigger overproduction of the reactive oxygen species (ROS) and cause DNA damage. This damage then cause adverse effects, such as abnormal transformation of skin cells and immunosuppression towards malignant cells. The typical phenotypic changes vary from benign changes such as wrinkles to fatal conditions such as blindness and the development of tumours and/or cancerous cells ^1 2^.

In response to UV radiation along with its DNA-damaging effects, the human skin employs endogenous mechanisms to prevent and/or cure the radiative effects. Examples of the mechanisms include apoptosis of mutated cells, overexpression of antioxidant genes, rapid repair of DNA, thickening of the epidermis and, most importantly, increased melanogen-esis ^3 4^. Melanogenesis, a biosynthetic pathway responsible for the production of melanin in the epidermal layer ^5^, is vital as the melanin produced as a result of the process, serves as a UVR absorbent, antioxidant and ROS scavenger ^3^. Thus increased melanogenesis is essential to photoprotection against the chronic effects of UV radiation including photo-aging, photodamage and photo-carcinogenesis, otherwise known as skin cancer. Especially post-exposure to UVR, melanin begins developing toxic properties. In-vitro studies testify to melanin’s ability to react and alter DNA negatively. Pheomelanin, additionally, has been evidenced to be especially vulnerable to photodegradation, generating hydrogen peroxide, superoxide anions, causing mutations in melanocytes and inducing apoptosis of cells among other dangers of increased melanin ^6^. Overaccumulation of melanin is also unfavourable as it might lead to a myriad of disorders, including, hyperpigmentation, neurodegeneration and even skin cancer ^7^. Tight regulation of skin melanin levels is thus crucial since both abnormally high and low melanin levels pose a potential danger for human well-being.

Regulation of melanin levels can be achieved indirectly through adjustments of tyrosinase activity. Tyrosinase is a rate-limiting enzyme responsible for the first step of melanin synthesis - conversion of tyrosine into L-3,4-dihydroxyphenylalanine (DOPA) ^5 7^. The other steps in melanogenesis are mostly spontaneous, therefore they require no catalyst. Due to these factors, overproduction of melanin is almost always due to overexpression of tyrosinase ^5 8^. The search for tyrosinase inhibitors is thus of utmost importance and has been an attractive aspect of research for the cosmetics and food industry as well as a central endeavour for the medical industry in inhibiting and minimizing the damaging effects of UVR.

Amongst the candidates for tyrosinase inhibitors are mushrooms. Mushrooms have been long known to be rich in bioactive agents ^9^, such as phenolic compounds. Additionally, mushrooms are appreciated for their antioxidant activity, primarily free radical scavenging effects, leading them to be valuable natural sources for pharmaceutical and nutritional purposes. Previous research, namely Gąsecka et al. ^7^‘s investigates the abundant composition of bioactive compounds and phenolic acids in seven strains of the Agaricus genus of mushrooms, one of which include the *Agaricus* bisporus white capped mushroom, evidencing the antioxidant and antiradical activities of this genus of mushrooms. While the sources from Chang et al. ^10^ and Boonsong et al. ^2^, do not study the specific Agaricus genus of mushrooms, they too evidence the value of mushrooms in antioxidant capabilities (mainly free radical scavenging, phenolic and flavonoid).

On melanin-reducing capabilities, the work of Zaidi et al. ^4^, having recognized a lack of research augmenting the claim that mushroom extracts can be effective as melanogenic agents, have investigated the “effect of purified mushroom tyrosinase of *Agaricus* bisporus on B16F10 melanocytes for the melanin production via blocking pigment cell machinery” and thus have testified that the mushroom *Agaricus bisporus* is in fact a viable agent for melanogenesis inhibition.

*Agaricus bisporus*, commonly known as the white button mushroom, are amongst the most conventional, cultivated gourmet species of mushrooms. It is one of the most popular edible mushrooms, native to European and North American grasslands. The mushroom belongs to the Agaricus genus of the Agaricacae family. The mushroom is recognized for its hemispherical white cap and stem, which flattens out through maturity, and is the young origin of the Portobello mushroom. A. bisporus contains various known beneficial nutrients such as vitamin D, potassium and sodium. Its consumption is also associated with a decreased risk of breast cancer and prevention of oxidative stress associated diseases^2 11^.

Despite prior works evidencing the potential for *Agaricus bisporus* to be natural sources of antioxidants and tyrosinase inhibitors, minimal work has been done to assess the potential of *Agaricus bisporus* as a preventive and curative agent against ultraviolet exposure. Thus in this work, we aim to correlate and understand the role of the secondary metabolites of *Agaricus bisporus* ethanolic extracts with regards with its antioxidant, sun-protecting-factor (SPF), and anti-tyrosinase inhibiting properties. The identification of secondary metabolites is done quantitatively with regards to total phenolic and total flavonoid contents, while other phytochemical screening is done qualitatively.

## Materials and Methods

### Sample Preparation

Cultivated gourmet mushroom species *Agaricus bisporus* is first collected and chopped into fine pieces. Then the finely chopped whole *Agaricus bisporus* was freeze dried for 72 hours. Freeze dried sample was ground using dry spice grinder into fine powder form.

### Ethanol Extraction

25 g of freeze-dried mushroom powder was extracted with 250 g of ethanol absolute. The maceration was left to sit for 24 hours at RT. The supernatant was filtered through Whatman filter paper in a vacuum pump. The recovered supernatants, after the filter process, were combined and the ethanol was removed by rotary evaporation. The obtained extract was stored in an amber sample vial, at 4 ^°^C.

### DPPH Radical Scavenging Assay

The DPPH assay aims to investigate the free radical scavenging activity of antioxidants (AH) towards the purple-coloured DPPH in et-OH, conducted as described by Lamien-Meda et al. methodology ^12^. Samples were prepared by serial-dilution method to yield 2, 4, 6, 8, and 10 ppm concentrations. Antioxidant assay is performed by adding 1 mL of 0.2 mM DPPH and 0.8mL of ethanol (et-OH). All the samples were prepared in triplicate, vortexed for 1 min and incubated in the dark for 30 min at room temperature. The absorbance of each sample and control was measured against 2mL of ethanolic DPPH as blank on UV–Vis Spectrophotometer (Hitachi U1800) at wavelength of 517nm. Percentage of DPPH inhibition was calculated using the formula (Equation 1), similar to the methodology used by Kumara et al. ^13^. The radical scavenging activity percentage (%RSA) can be calculated from Equation 1.

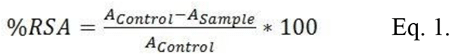

where A_*Control*_ is the absorbance of 0.2 mM ethanolic DPPH only and A_*Sample*_ is the absorbance of the reacting mixture, with the addition of the mushroom sample. IC50 value (concentration at 50% inhibition) was obtained from the linear regression of the data points and extracting the concentration at 50% inhibition value.

### Sun Protecting Factor (SPF) Test

Methodology used is as described in Romulo et al.’s ^14^. The absorption characteristics of the sunscreen agents against UVB (290-320 nm) can be determined in vitro by utilizing UV spectrophotometry with the following Equation 2 below:

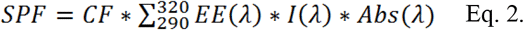

where: EE (l) – erythemal effect spectrum; I (l) – solar intensity spectrum; Abs (l)-absorbance of sunscreen product ; CF – correction factor (= 10). The values of EE x I at different wavelengths are given in Table 1 according to work done by Romulo et al’s. ^14^.

**Table 1.** EExI (normalized) at each wavelength. ^14^.

| Wavelength (nm) | EE x I (normalized) |
| --- | --- |
| 290 | 0.0150 |
| 295 | 0.0817 |
| 300 | 0.2874 |
| 305 | 0.3278 |
| 310 | 0.1864 |
| 315 | 0.0839 |
| 320 | 0.0180 |
| Total | 1 |

Sample was dissolved in ethanol to a concentration of 5,000 ppm. The absorbance of the sample solution was then read at wavelengths 290 - 400 nm with 5 nm interval. Ethanol was used as the blank, negative control. To obtain the SPF value, the absorbances of the sample at various wavelengths listed in Table 1 are obtained and then plugged into Equation 2. Their summation gives the total SPF value.

### Tyrosinase Inhibition

Tyrosinase inhibition activity was determined according to the modified method of Chan et al, ^15^ or Burger et al. ^16^. Sample was dissolved in dimethyl sulfoxide (DMSO) to a concentration of 10,000 ppm and then diluted in potassium phosphate buffer (50 mM, pH 6.8) to 0.05 ppm. All steps were conducted at room temperature. Samples with different concentration (70 *μ*l) was incubated with 30 *μ*l of the tyrosinase enzyme (333 U/ml in phosphate buffer, pH 6.8) for 5 min, followed by the addition of 2 mM of the substrate, L-tyrosine (110 *μ*l) and further incubated for 30 min at RT, ensuring that effect of light is limited. Kojic acid was used as positive control. The mixture of sample and other components, except L-tyrosine, was used as a blank. Absorbance reading was performed at 475 nm. The percentage of tyrosinase inhibition was calculated using Equation 3:

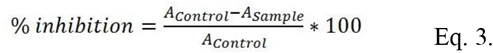

where, %inhibition is the tyrosinase inhibition percentage, A_*control*_ is the absorbance of the control solution and A_*sample*_ is the absorbance of the sample solution. For each sample concentration, the absorbance was recorded in triplicate manner.

### Total Phenolic Content by Folin Reagant Method

Total phenolic content in A. bisporus extracts was determined using the Folin-Ciocalteu reagent using the method of Javanmardi et al. ^17^. Serial dilution of gallic acid stock solution was made with the concentration of 100 ppm, 80 ppm, 60 ppm, 40 ppm and 20 ppm. For Folin-Ciocalteu stock solution, 10 mL of Folin reagent was diluted into 100 mL (10% w/v) of distilled water and then stored away from light. Na_2_CO_3_ (Sodium Carbonate) solution was prepared by diluting 7.5% (w/v) of Na_2_CO_3_ in distilled water solvent. Standard curve was prepared by combining 1.5 mL of the Folin-Ciocalteu reagent and 0.3 mL of gallic acid into a test tube. The mixture was vortexed and incubated in the dark for 5 minutes at room temperature. After incubation, 1.2 mL of Na_2_CO_3_ was added and vortexed once more. The solutions were then incubated in the dark for 15 minutes. The absorbance of each sample was measured against 0.3 mL distilled water + 1.5 mL Folin reagent, using 1.2 mL Na_2_CO_3_ as blank solution, and measured by UV–Vis Spectrophotometer (Hitachi U1800) at a 765 nm wavelength. Gallic acid was used as the standard curve. Sample results were presented in mg gallic acid equivalent (GAE)/mL.

### Total Flavonoid Content

The method is adopted from Mansur et al. ^18^. The standard curve determination was carried out using 5, 10, 15, 20, 25, 30, 35, 50 ppm serial dilution of quercetin in Et-OH. 2 grams of aluminum chloride was diluted in 100 mL Et-OH. The experiment is conducted in a fume hood as AlCl_3_ is corrosive. 2 mL of AlCl_3_ and 2 mL of each quercetin concentration solution were transferred into individual test tubes, incubated, vortexed and measured at 415 nm using a spectrophotometer. For sample measurement, 2 mL of the sample ethanolic extract is added to 2 mL AlCl_3_ (2% solution) in each tube and then measured. The sample is presented in mg quercetin equivalent (QE)/mL.

### Qualitative Phytochemical Screening

#### Alkaloids

Identification of Alkaloids was done via methods described in Farnsworth, N. ^19^. and Depkes RI ^20^. The sample contains alkaloids if at least two groups of the experimental solutions from the three reagents tested form precipitates. To prepare the test solution, 50 mg of the extract was added to 1 mL HCL 2 N and 9 mL water and then it was heated in a water bath for 2 minutes, cooled and filtered to obtain the filtrate.

#### Bouchardat’s Reagent

The reagent was made of 2 g iodium P and 4g Kl P that were dissolved in 100.0 mL distilled water. 1mL filtrate was added to 2 drops of the Bouchardat Reagent. If a brown-black precipitate is formed, the sample is positive, containing alkaloids.

#### Mayer’s Reagent

The reagent was made of a mixture of a solution of mercury (II) chloride P (1.358 g of HgCl_2_ in 60 mL of distilled water) and a solution of potassium iodide P (5 g of potassium iodide in 10 mL of distilled water) was combined with distilled water until the volume of the entire mixture reached 100 mL. 1 mL filtrate was added to 2 drops of the Mayer’s Reagent. If a lumpy white or yellow precipitate is formed that dissolves in methanol, the sample is positive, indicating the presence of alkaloids.

#### Dragendorff’s Reagent

The reagent was made of a mixture of bismuth nitrate P (8g of bismuth nitrate in 20mL nitric acid) and potassium iodide P solution (27.2 g potassium iodide P in 50.0 mL distilled water) which was left to stand until completely separated. The clear solution is taken and dissolved in distilled water until the entire mixture’s volume reaches 100.0 mL. 1 mL of the filtrate was added to 2 drops of the Dragendorff’s Reagent. If an orange-brown precipitate is formed, the sample is positive, containing alkaloids.

#### Flavonoid

Identification of flavonoid content was done by following the procedure in ^20^. A few mg of extract was added to 4mL ethanol P until the extract was dissolved. 2 mL of the solution was added to 0.5 grams of zinc powder and 2 mL of 2N HCl and was left to stand for 1 minute. Then 10 drops of concentrated HCl P were added. An intensive red colour observed within 2-5 minutes indicates the presence of flavonoids (glycoside-3-flavonol).

#### Tannin

Identification of Tannin was done as illustrated in Farnsworth, Norman R.’s work ^19^. A few mg of the thick extract was added to 15 mL of hot water. Then heated until mixture boils for 5 minutes. Filtrate was filtered. A few drops of 1% FeCl_3_ was added to produce a green-violet colour.

#### Saponin

5 mg of the extract was added to a test tube to which 10 mL of hot water was also added. Mixture was cooled and shaken vigorously for 10 seconds. A steady foam formed 1 to 10cm high for no less than 10 minutes. With the addition of 1 drop of 2N HCl, the foam did not disappear.

#### Terpenoid

A total of 5 mg of the concentrated extract was added to 3 mL of dichloromethane and then evaporated in an evaporating dish. The evaporated residue is added to 6 drops of acetic acid and 3 drops of concentrated H_2_SO_4_. The sample is positive for terpenoids if it produces a red-green or violet-blue colour.

## Results and Discussion

As has been structured in the preceding portion of the article, results are reported in two parts, that of the assessments of the properties of A bisporus mushrooms and secondly, identification of the compounds responsible for the observed abilities of the A bisporus.

### Part 1: Assessment of Antioxidant, SPF and Tyrosinase-Inhibiting Properties

#### DPPH Radical Scavenging Activity

The *Agaricus bisporus* extract’s free-radical scavenging capacity was measured by the DPPH (2, 2-diphenyl-1-picrylhydrazyl) method. The assay relies on measuring the absorbance of the naturally purple-coloured DPPH with respect to how much the sample can quench it, rendering the solution colourless. The antioxidant compound donates a hydrogen atom to the DPPH radical solution which is then reduced. The absorption of the hydrogen by the antioxidant, namely the A bisporus given that it contains the bioactive compounds for the mushroom to act as an antioxidant, is in stoichiometric ratios with respect to the level of reduction and the DPPH remaining after the radical-scavenging effect. Upon measurement following a set amount of time, the absorption/reduction of the hydrogen is inversely related with the radical scavenging capabilities of the antioxidant ^21^. The results were reported as IC_50_ values, the concentration required to achieve 50% RSA.

As a baseline for understanding the *Agaricus bisporus*’s ability to scavenge radicals compared to a given standard, a brief review of existing literature will be presented prior to discussion of the results. Abdullah et al.’s ^21^, research of selected culinary medical mushroom’s antioxidant effects reports positive control (quercetin) DPPH scavenging ability of IC50 = 0.03 mg/mL whose value can be interpreted as excellent, while the various mushroom extracts exhibited radical scavenging ability of 5.28-35.6 mg/mL which was suggested still to be viable (although quite low in general) for prospects of the mushroom sample being primary antioxidants whose consumption could reduce the risk of cardiovascular disease and act as a protective agent against oxidative stress.

Another study by Gasecka, et al. ^7^ reports an EC_50_ value for DPPH radical scavenging effective concentration, that ranges from ∼800 to 3200 *μ*g/mL for the A.bisporus species. Amongst the mushroom species investigated in Gąsecka, et al.’s study, the white A.bisporus showed the least effective inhibition.

As a comparison of what is considered a highly effective antioxidant, in Table 2, results of Di Petrillo et al.’s ^22^ study is shown, elucidating the antioxidant properties of the derived ethanolic extracts of Asphodelus microcarpus medicinal plants, with DPPH scavenging IC50 values of 55.9 ± 1.55 for the leaves of the plant, 28.4 ± 0.85 for its flowers and 360 56.57 for its tubers.

**Table 2.** Antioxidant activity of *Agaricus bisporus*. (DPPH) radical scavenging assay.

| Concentration | Abs | %RSA |
| --- | --- | --- |
| Control | 1.249 | - |
| 10000 | 0.196 | 84.307 |
| 8000 | 0.310 | 75.180 |
| 6000 | 0.550 | 55.965 |
| 4000 | 0.752 | 39.792 |
| 2000 | 1.034 | 17.214 |

In Table 2 and Figure 1, the %RSA of the A bisporus ethanolic extract is shown. Results conclude that the A bisporus ethanolic extract presents IC50 of ∼ 5456 *μ*g/mL. In Table 3, a summary of IC50 values from previously reported works are compiled and used for result comparison. A lower IC50 value is indicative of stronger antioxidant activity, hence, using Abdullah et al.’s ^21^ work as a basis for comparison, the ethanolic extract of A.bisporus mushrooms is considered moderately efficient as radical scavenging agents. In Abdullah et al.’s work ^21^, a much higher IC50 is reported. However it must be noted that in the mentioned study, the group evaluates the possibility of using hot water for extraction as this would yield the least amount of chemical residues in the product. The high temperatures used in the extraction may cause detrimental effects to the metabolites in the samples, thus leading to a very high IC50 value.

**Table 3.** Comparison of DPPH radical scavenging ability of *Agaricus bisporus* ^7^ other mushrooms ^21^ and *Asphodelus microcarpus* ethanolic extract ^22^.

| | *Asphodelus microcarpus* (ethanolic extract, $\mu\text{g/mL}$)<sup>22</sup> | | | Various mushroom species (hot water extract, $\mu\text{g/mL}$)<sup>21</sup> | *Agaricus bisporus* various strain (70% ethanol extract, $\mu\text{g/mL}$)<sup>7</sup> | *Agaricus bisporus* (ethanol extract, $\mu\text{g/mL}$) [This Work] |
| --- | --- | --- | --- | --- | --- | --- |
|  | Leaves | Flowers | Tubers |  |  |  |
| DPPH scavenging IC50 | ~55.9 | ~28.4 | ~360 | ~5280-35659 | ~800-3200 | ~5456 |

**Figure 1.**
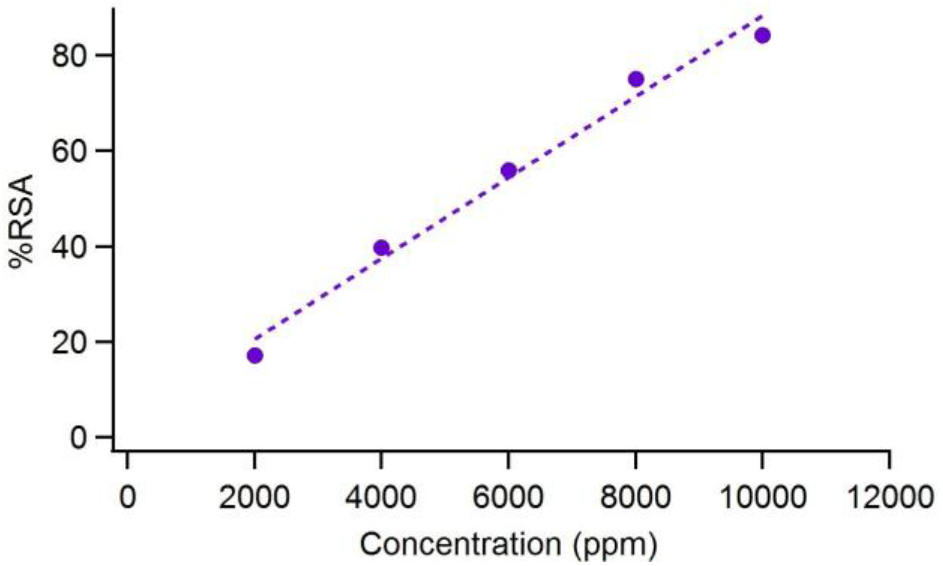
Antioxidant activity of *Agaricus bisporus* ethanolic extract.

Comparing with the results reported by Gąsecka, et, al. ^7^, we find that our IC50 value is slightly higher than theirs. This difference may stem from the fact that Gasecka et al. utilizes 70% ethanol as extraction solvent whereas in our work absolute ethanol is used. The diluted ethanol may be more effective in extracting the metabolites that are responsible for the antioxidant property, in line with similar observations in other studies ^23^.

Though Di Petrillo’s ^22^ research is not of any species of mushrooms, rather a medicinal plant with vastly differing features than the A. bisporus mushrooms, it is of notable significance to note that compared to the range of IC50 values in *μ*g/mL observed for the plant, from 28.4 in the flowers to 360 in the tubers.

### SPF Test

A very minimal amount of earlier works have pioneered the research of SPF of specifically the A.bisporus mushroom species. Suhaenah et al.’s ^24^ work amongst one of the scanty available pieces of literature, reported a range of SPF values at different concentrations, of 17.672 up to 31.325 at higher concentrations, evidencing the claim that the A.bisporus mushroom species is capable to providing medium to the lower end of high SPF, protection against UVR.

Additionally, Hailun et al’s ^25^ comprehensive review of natural components commonly found in market sunscreens report various compounds such as Silymarin, an essential compound for topical photoprotection, renowned for its antioxidant properties and potential as a protector against solar radiation, independently having an SPF value of 5.50 0.25, at a concentration of 50 *μ*mol/L, processed as an ethanolic solution and analysed in vitro. When combined with chemical sun-protecting agents titanium dioxide and zinc oxide, the observed effect was an increased SPF ability, at 12.37 ± 4.39 and 16.30 ± 2.98, respectively. Plant-derived extracts that have been recently incorporated in market sunscreens including Sphaeranthus indicus, rich in phenols and flavonoids were found to have SPF values of 3.85 at 10000 ppm^26^. The renowned cancer-preventing plant Moringa oleifera, rich in polyphenols, some of which include “quercetin, rutin, chlorogenic acid, ellagic acid, and ferulic acid that can be applied in sunscreens” report SPF values of 2 at concentration of 20000 to 40000 ppm^27^. Chitosan/tripolyphosphate (TPP) nanoparticles “with vegetable extracts rich in flavonoids provides a topical formulation against sunburns” and were reported to have an SPF of 2.3 ± 0.4 at 200 ppm concentration ^28^. Furthermore, the study highlights the synergic effects of these natural ingredients and extracts in market sunscreen formulation with many of the natural sunscreens enhancing the protective effects of chemical sunscreens.

In our work we find that the A.bisporus ethanolic extract with a concentration of 5,000 ppm has SPF value of 5.355 (Table 4). This value is relatively low in comparison to A. bisporus ethanolic extract reported by Suhaenah et al. ^24^ Based on their reported data for extract with 1% to 4% solution, which equates to 10000 to 40000 ppm, by extrapolating the dataset (Figure 2), we can expect an SPF value of ∼11 for their samples at 5000 ppm (Figure 2). This is higher than the SPF value found in our work. In comparison with other compounds, the general standard for low, medium and high SPF values are between 2 to 11, 12 to 29, and 30 to 50, respectively ^24^. The higher the concentration, the higher the SPF value in general. In most cases, concentration of extract under test ranges between 0.1 to 10% or 1000 to 100000 ppm.

**Table 4.** SPF of *Agaricus bisporus* ethanolic extract at 5000 ppm concentration. Raw data calculations from absorbance at wavelengths 290-320, sample diluted with dilution factor 3.

|  | Abs | Dilution factor | Abs Final | EE x I | Abs x EE x I | Total | Correction factor | SPF |
| --- | --- | --- | --- | --- | --- | --- | --- | --- |
| 290 | 0.46 | 3 | 1.386 | 0.0150 | 0.02 |  |  |  |
| 295 | 0.43 | 3 | 1.293 | 0.0817 | 0.11 |  |  |  |
| 300 | 0.23 | 3 | 0.678 | 0.2874 | 0.19 |  |  |  |
| 305 | 0.13 | 3 | 0.390 | 0.3278 | 0.13 | 0.536 | 10 | 5.355 |
| 310 | 0.10 | 3 | 0.312 | 0.1864 | 0.06 |  |  |  |
| 315 | 0.09 | 3 | 0.282 | 0.0839 | 0.02 |  |  |  |
| 320 | 0.09 | 3 | 0.255 | 0.0180 | 0.00 |  |  |  |

**Figure 2.**
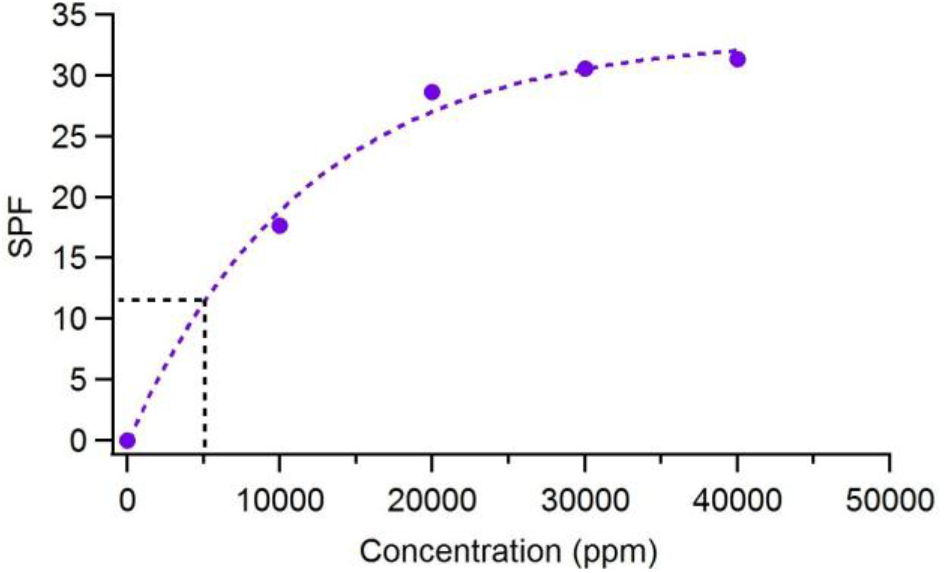
Extrapolation of SPF vs concentration data reproduced from ^24^ regarding ethanolic extract of *Agaricus bisporus*. Based on this extrapolated data, SPF of ∼ 11 is expected for 5000 ppm in the samples of Suhaenah et al. ^24^

Although the standard classification for SPF strength given by Suhaenah et al. ^24^ and regarded universally with regards to conventional sunscreens would categorize an extract with 5000 ppm concentration having the value 5.355 as being a low SPF agent, when viewed in the context of it being a natural ingredient, this value is relatively favorable. This is bolstered by the similar and even lower values reported in Hailun et al’s ^25^ study for the various natural compounds reviewed in their study such as the Sphaeranthus indicus, Moringa oleifera, Ginkgo biloba L., and others. Furthermore, it is interesting to note Hailun et al’s study’s ^25^ remarks on the importance of viewing natural agents as enhancers rather than stand-alone UVR protectors. A.bisporus mushrooms may likewise pose as valuable agents to be further studied and utilized in the skin-care and medical industry with regards to the excessive melanogenesis discussed earlier.

### Tyrosinase Inhibition Test

Tyrosinase, as aforementioned, is an oxidase comprised of copper and an enzyme that catalyses the synthesis of melanin. The rate-limiting enzyme has the role of converting tyrosine into dihydrophenylalanine (DOPA) and further oxidize this DOPA into dopaquinone. Melanin inherently is a natural mechanism of the human body however the over-secretion of melanin from melanocytes is what comprises the main health issue, caused by external factors such as exposure to UVR and release of *α*-melanocyte-stimulating hormones ^29^. The overexpression of tyrosinase therefore leads to the overproduction of melanin associated with hyperpigmentation, freckles or melanoma. Hence, tyrosinase inhibitors have attracted much attention by various industries, not only the cosmetic industry but also in medicine and pharmacology. Kojic acid, a synthetic tyrosinase inhibitor widely utilized in skin-whitening agents, is used as a positive control in the current study. However, these synthetic compounds have been evidenced to be associated with side effects such as erythema or contact dermatitis, increasing interest towards natural and biocompatible agents ^30^, sparking the interest of determining the tyrosinase inhibiting activities of A.bisporus mushrooms, whose properties with regards to tyrosinase inhibition is minimally documented in literature.

As a baseline for evaluation, Di Petrillo et al.’s ^22^ results are tabulated in Table 5 showing tyrosinase inhibiting ability of Asphodelus microcarpus medicinal plants with an IC50 value of 270 *μ*g/mL. Furthermore, the tyrosinase inhibiting properties of A.bisporus and other mushrooms studied by Taofiq et al. ^29^ are also listed for comparison. They ^29^ found that A.bisporus ethanolic extracts had anti-tyrosinase potential with an EC50 value in mg/mL of 160 *μ*g/mL, the highest among the studied mushroom species which included P.ostreatus and L.edodes. In addition, the literature cited in Taofiq et al.’s report, namely Miyake et al.’s ^31^ confirms that the phenolic compound 2-amino-3H-phenoxazin-2-one, isolated from A.bisporus, at 0.5 *μ*M, could significantly inhibit the enzyme tyrosine up to 80%. To further reinforce this, Yan et al. ‘s study ^32^ describes that the isolated phenolic acids: steroidal triterpenes, namely botulin and trametenolic acid, found in mushroom extracts inhibit tyrosinase activity with an IC50 of 5.13 *μ*M, stronger than the IC50 value of their positive control kojic acid at 6.43 *μ*M, and 7.25 *μ*M respectively. These results suggest that the constituent bioactive phenolic compounds of mushrooms, including A.bisporus can be exploited as tyrosinase-inhibiting agents. Their work is consistent with that of Di Petrillo et al ^22^ on the phenolic rich Asphodelus microcarpus which is shown to exhibit high antioxidant and tyrosinase inhibition properties.

**Table 5.** Comparison of tyrosinase Inhibition *of Agaricus bisporus* in this work against those in ^29^ and *Asphodelus microcarpus* ethanolic extract reported in ^22^.

| | *Asphodelus microcarpus* ($\mu\text{g/mL}$) <sup>22</sup> | | | *Lentinula edodes* ($\mu\text{g/mL}$) <sup>29</sup> | *Postreatus ostreatus* ($\mu\text{g/mL}$) <sup>29</sup> | *Agaricus bisporus* ($\mu\text{g/mL}$) <sup>29</sup> | *Agaricus bisporus* ($\mu\text{g/mL}$) [This Work] |
| --- | --- | --- | --- | --- | --- | --- | --- |
|  | Leaves | Flowers | Tubers |  |  |  |  |
| Tyrosinase Inhibition IC50 |  | ~270 |  | ~820 | ~860 | ~160 | ~2 |

In our work, we find that the *Agaricus bisporus* ethanolic extract exhibits an exceptionally high tyrosinase inhibiting activity. As compared to kojic acid as the positive control, the *Agaricus bisporus* ethanolic extract shows significantly higher inhibition of tyrosinase (Figure 3). IC50 value of kojic acid was 135.50 ppm or *μ*g/mL. On the other hand, the extract sample shows strong activity at very small concentration. Almost complete inhibition is observed for samples with concentration larger than 3 *μ*g/mL. Between 0.01 *μ*g/mL and 3 *μ*g/mL, we observed a gradual increase in the tyrosinase inhibition. Figure 3a shows the data for three separate trials, showing similar trends. It was quite challenging in obtaining precise IC50 for the extract due to the very low concentration of extract. From Figure 3.a, IC50 for the *Agaricus bisporus* ethanolic extract is estimated to be ∼ 2 *μ*g/mL. Our result shows stronger tyrosinase inhibition capability than that reported by Taofiq et al. ^29^. In general, our result is consistent in that there is a substantial tyrosinase inhibiting properties in the *A*.*bisporus* mushroom as reported by Taofiq et al. ^29^, Miyake et al. ^31^ and Yan et al. ^32^. The A.bisporus ethanolic extract in our work is also shown to have better tyrosinase inihibiting performance compared to that by Asphoedulus microcarpus in Table 5^22^, further supporting the conclusion that A.bisporus mushrooms are superior tyrosinase inhibitors and thus hyperpigmentation or other melanin-related disorder correctors.

**Figure 3.**
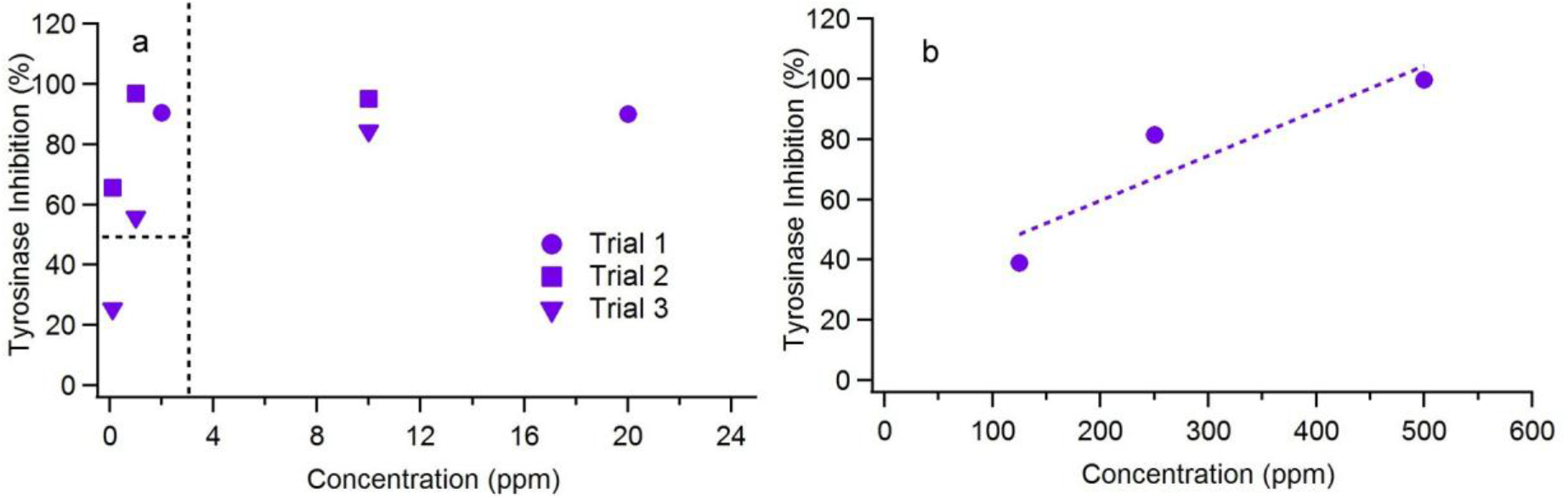
Tyrosinase inhibition of (a) *Agaricus bisporus* ethanolic extract in this work in comparison to the (b) kojic acid control samples. Dotted lines in (a) are guiding lines.

The difference between our values with that of Taofiq et al ^29^ might be associated with the slightly different extraction methods. In their work, samples are first dried at 30°C in an oven and processed into fine powder, followed by 12 cycles of reflux for 4 hours in an assumed 78 °C (ethanol reflux temperature), and finally evaporated under reduced pressure using a rotary evaporator to obtain the dried ethanolic extract. In our work, relatively milder sample processing conditions are used which may slightly improve the preservation of the metabolites in the sample. Samples are first freeze fried for 72 hours and ground into fine powder. Maceration was done at room temperature for 24 hours followed by filtration and solvent evaporation under reduced pressure in a rotary evaporator.

### Part 2: Identification of Phenolic, Flavonoid, and Other Phytochemicals

To gain insight on the possible origin of the observed properties of the *Agaricus bisporus* ethanolic extract, next we report selected phytochemical analysis that we carry out. Total phenolic content analysis is carried out quantitatively while the flavonoid content analysis is carried out both quantitatively and qualitatively. Other secondary metabolites are screened qualitatively.

### Total Phenolic Content by Folin Reagant Method

Phenolic compounds are commonly found in many plants, namely also mushrooms and are indicative of and closely related with antioxidant activities.

Gąsecka, et, al. ^7^ studied the profile of phenolic, organic acids and antioxidant properties of various species of Agaricus mushrooms. Their report demonstrated that various strains of white A. bisporus mushroom species has a total phenolic content in the range of ∼ 132-756 mg GAE/100g DW.

Furthermore, Alispahic’ et al.’s ^33^ study of the phenolic content of various species of mushrooms (dry boletus, white and brown champignon, oyster and shiitake mushrooms) showed total phenolic content of ∼ 494-3556 mg GAE/100gand concluded that these mushroom species were potential sources of natural antioxidants. The highest total phenolic content was observed in dry boletus mushrooms.

In this work, we determine the total phenolic content in *Agaricus bisporus* ethanolic extracts to be ∼1143 mg GAE/100g (Figure 4 and Table 6). This value exceeds that of Gąsecka, et, al.’s ^7^ and lies within the range of values reported by Alispahic’ et al ^33^. The different total phenolic content compared to previously reported work could be due to different extraction methods.

**Table 6.** Total phenolic content of *Agaricus bisporus* ethanolic extract measurement and calculation.

|  | Abs. | Conc. (mg/L) | Unit Conv. [x 1L/5g (mg GAE/g)] | Ave. (mg GAE/100g) |
| --- | --- | --- | --- | --- |
| Sample 1 | 0.562 | 49.33945 | 9.878 | 1143.2 |
| Sample 2 | 0.732 | 64.93578 | 12.987 |  |

**Figure 4.**
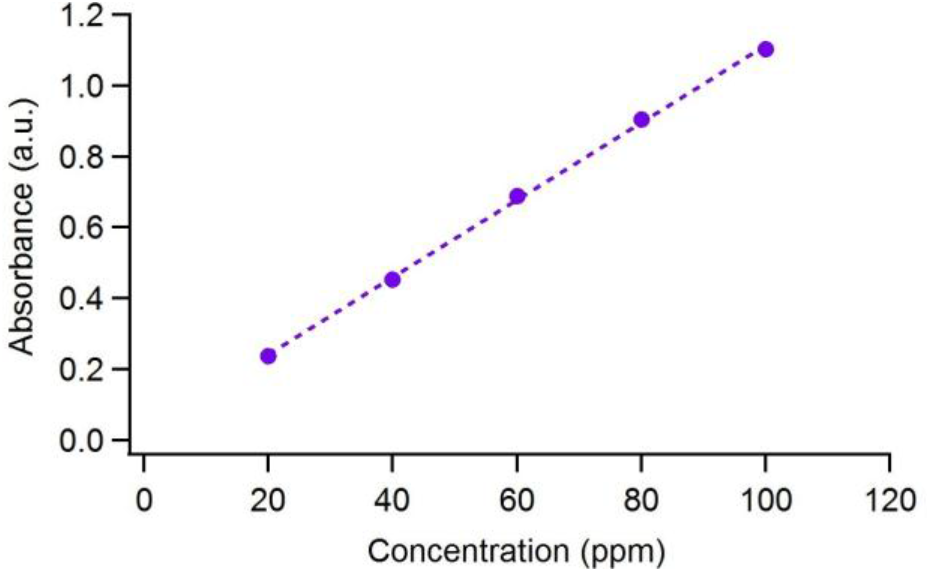
Calibration curve of gallic acid standard for determination of total phenolic content of *Agaricus bisporus* ethanolic extract.

### Total Flavonoid Content

Flavonoids are a group of natural substances with variable phenolic ^34^, generally found in vascular plants as glycosides and flavonoid aglycones. Typically in analysing flavonoids, the aglycones in the plant extracts that have been hydrolysed are examined. The flavonoid extraction is carried out with boiled ethanol to avoid enzyme oxidation ^35^. Flavonoids are renowned for a broad spectrum of favourable health-promoting benefits and hence are considered indispensable components in diverse applications in the nutraceutical, pharmaceutical, medicinal and cosmetic industry. This can be attributed to their “anti-oxidative, anti-inflammatory, anti-mutagenic and anti-carcinogenic properties coupled with their capacity to modulate key cellular enzyme function” with an added impulse for lowering cardiovascular mortality rate and association with diseases such as cancer, Alzheimer’s disease and atherosclerosis among others ^34^.

Gil-Ramírez et al.’s ^36^ study which testifies to the claim that mushrooms do not contain flavonoids and cannot actually synthesize flavonoids as they lack the required enzymes. The general trend in scientific literature testifying to the presence of flavonoids in mushrooms is critiqued for their uses of unspecific colorimetric methods developed originally to quantify flavonoids in plants as a general category. Furthermore, sequences of DNA encoding for the key enzymes involved in the flavonoid biosynthetic pathway were not discovered in the genomic databases of completely sequenced mushrooms. It was evidenced that the fruiting bodies of mushrooms were also incapable of absorbing flavonoids from their surroundings. These results also agree with that of Gąsecka, et, al.’s ^7^ study that reports that flavonoids were not found in all the mushroom species experimented, including white *Agaricus bisporus*.

The determination of the total flavonoid content in *Agaricus bisporus* was carried out with ethanol extracted samples using quercetin as standard (Figure 5 and Table 7). Total flavonoid content of A.bisporus mushrooms is 0.759 mgQE/g, a very minimal quantity. In retrospect with past studies, these results agree with that of Gil-Ramírez et al.’s ^36^ and Gąsecka, et al.’s ^7^, confirming the absence or very minor amount of flavonoids in *Agaricus bisporus* mushrooms. The lack of flavonoid in our sample is also confirmed by the qualitative phytochemical screening shown in Figure 6.

**Table 7.** Total flavonoid content of *Agaricus bisporus* ethanolic extract measurement and calculation.

|  | Abs. | Conc. (mg/L) | Unit Conv. [x 1L/10g (mg QE/g)] | Ave. (mg QE/g) |
| --- | --- | --- | --- | --- |
| Sample 1 | 0.206 | 8.072 | 0.807 | 0.759 |
| Sample 2 | 0.178 | 7.113 | 0.711 |  |

**Figure 5.**
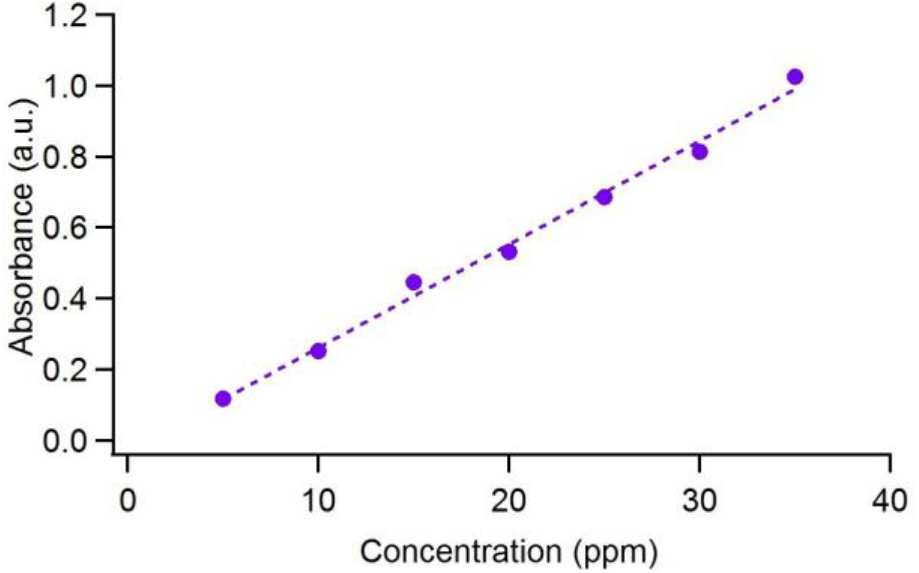
Calibration curve of quercetin standard for determination of total phenolic content of *Agaricus bisporus* ethanolic extract.

**Figure 6.**
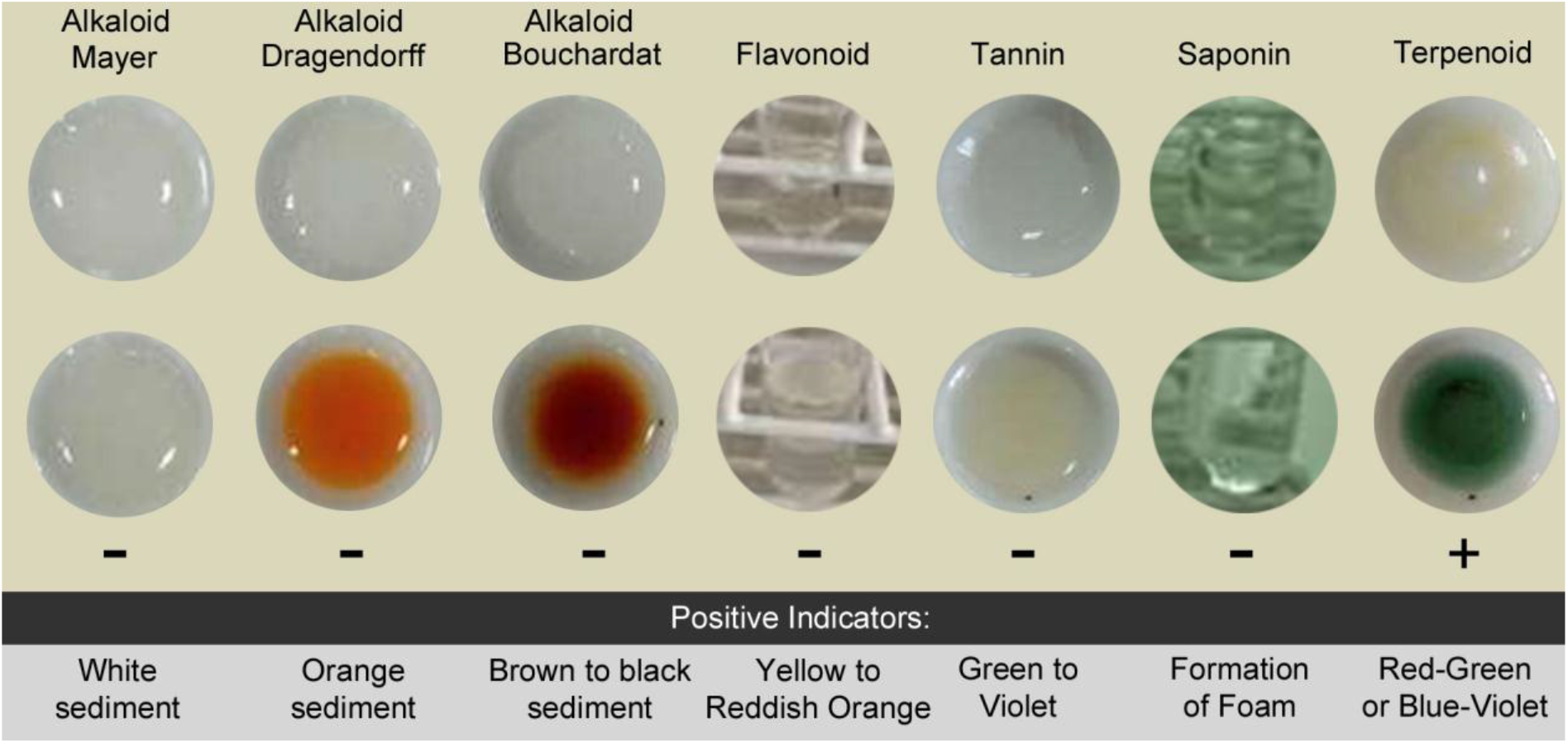
Qualitative phytochemical testing observation. Only terpenoid is clearly detected in the *Agaricus bisporus* ethanolic extract.

### Qualitative Phytochemical Screening

The phytochemical screening is a qualitative test of the chemical contents of plants in order to determine the class of compounds contained in the extracts derived from plants. The screening analyses the presence of secondary metabolites like alkaloids, flavonoids, terpenes, tannins, saponins, terpenoids, glycosides, quinones and anthraquinones in the extracts^35^.

Several works reported phytochemical screening of *Agaricus bisporus*. Eswari et al. ^37^ studied A. bisporus petroleum ether and methanolic extracts and found that alkaloid is only detected in the petroleum ether based extracts. Flavonoid was weakly detected in the methanolic extract and not in the petroleum ether extract. Tannin was strongly detected in petroleum ether extracts and not in the methanolic extract. Saponins are strongly detected in methanolic extract and not in petroleum ether extract. Terpenoid is weakly and moderately detected in the petroleum ether and methanolic extracts, respectively.

In our works, the extraction was carried out with absolute ethanol. The ethanolic extract in our work shows only the presence of terpenoid (Figure 6). This result generally agrees with the work of Eswari et al ^37^ except for the lack of detection for saponin. The presence of terpenoid indicates that A. bisporus has another potential application in the field of therapeutics medicine. Indeed, there appears to be renewed interest in the study of mushrooms as a natural source of terpenoids ^38^. Several groups have reported the efficacy of terpenoids for therapeutic treatments of neurodegenerative ^39^, antiviral ^40^, and more. Several studies have looked at more closely the health benefits of terpenoid extracted from various mushrooms such as Ganodherma ludicum ^41^, Antrodia cinnamomea ^42^, Inonotus obliquus ^43^, and others, but to the author’s knowledge, there has not been a study of therapeutics effects of terpenoid extracts from *Agaricus bisporus*. The presence of terpenoid could also be one of the high tyrosinase inhibition activity in the *Agaricus bisporus* in our work. Both phenolic and terpenoid compounds have been found to directly contribute to tyrosinase inhibiting activities ^44^.

Several things are interesting to note. First, our work agrees with that of others in that there is an absence or very minimal amount of flavonoid in the *Agaricus bisporus* mushrooms. Our work supports the prior theory and analysis which suggest the inability of the mushroom to produce flavonoids due to the lack of required enzyme and/or that the mushroom fruiting body is incapable of absorbing flavonoids from the surrounding.

Secondly, our work agrees with the prior reports ^7, 21^ that demonstrated *Agaricus bisporus* as the least effective radical scavenging agents compared to other common edible mushrooms. It is interesting also to note that our sample exhibits a higher amount of total phenolic content compared to that of *Agaricus bisporus* in other works ^7, 21^ but we observed a lower radical scavenging capability in our sample than expected. Thus, it can be said that the phenolic compound in *Agaricus bisporus* plays a minor role in radical scavenging.

Likewise, a lower value of SPF is also found compared to that of other works ^25^ but remains favorable compared to some plant sources ^24^. It is unclear whether the total phenolic content in our *Agaricus bisporus* is higher or lower than that in ^25^ since it was not reported.

The high amount of total phenolic content in our sample does however correlate with the exceptional tyrosinase inhibition capability we are seeing in our *Agaricus bisporus* sample. Such correlation between phenolic and tyrosinase inhibition have been seen in many systems ^38^. The correlation is further bolstered by the fact that our qualitative phytochemical screening established no alkaloid contents in the *Agaricus bisporus* ethanolic extract we examine. It is known that alkaloid is one of the drivers for tyrosinase inhibition activity ^38^.

Besides alkaloid, terpenoid is also known to be one of the drivers for tyrosinase inhibition activity ^38^ and indeed the terpenoid is present in our sample. It is not known how much is present in the sample but it is plausible that terpenoid in the sample contributes to both the radical scavenging and tyrosinase inhibiting properties we observe in our sample.

Thus, from the aforementioned results and discussion, it is observed that the strong tyrosinase inhibition in the *Agaricus bisporus* ethanolic extract is likely to be driven by the phenolics and terpenoids in the sample. More work however is required to elucidate further the types and distribution of phenolics and terpenoids in the sample among other more quantitative phytochemical analysis. We note that in our work we have minimized the use of higher temperature processings during the extraction and that we are using absolute ethanol as solvents instead of the more commonly used methanol and diluted ethanol. This combination of processing may result in different metabolites being extracted, which gives rise to some different behavior compared to other *Agaricus bisporus* work.

## Conclusion

In conclusion, our work agrees with previously reported work with some variations. We find that indeed the common button mushroom namely the *Agaricus bisporus* is available as a minor source of antioxidant agent with %RSA IC50 of ∼5456 *μ*g/mL. It also presents as a moderate source of sun protecting agent with SPF value of ∼5.355 at 5000 ppm concentration, prior to combination and supplementation with chemical UV filters such as titanium oxide or zinc. More work is required to further evaluate the SPF in our sample. It would be especially useful to obtain more SPF value at higher concentrations in order to compare its efficacy with those in other works. Of note, we observed a high tyrosinase inhibiting capability of the *Agaricus bisporus* ethanolic extract with IC50 of ∼2 *μ*g/mL. This is seen to be likely attributable to the phenolic contained in the mushrooms, found to have a value of ∼1143 mg GAE/100g and possibly partially attributable to the terpenoid present in our sample. Additionally, this work highlights the complex and sensitive biochemistry present in the seemingly common mushroom where it is possible that slight differences in extraction method, geographical conditions and cultivation would result in different properties of the extracted products. Overall, through this assessment, it is clear that *Agaricus bisporus* presents a really exciting outlook for the cosmetic industry and the field of pharmaceuticals where the search for natural compounds over synthetic products has been burgeoning with the rising concerns and prevalent health-issues related with UV radiation and exposure to free-radicals being the cause of oxidative stress.

## Acknowledgement

We are grateful for the funding and support of Sekolah Pelita Harapan Lippo Village through its Applied Science Academy program. The authors are thankful for the generous support and access to the research facility at Universitas Pelita Harapan and Emmerich Research Center. C.Y.H., Y.H., M.S., D.R. designed the experiment and carried out the extraction, purification, and characterization of the materials. C.Y.H., Y.H., S.P.W., E.S. carried out the analysis and wrote the manuscript.

